# Single-cell evidence for plasmid addiction mediated by toxin-antitoxin systems

**DOI:** 10.1101/2023.08.29.555305

**Authors:** Nathan Fraikin, Laurence Van Melderen

## Abstract

Toxin-antitoxin (TA) systems are small selfish genetic modules that increase vertical stability of their replicons. They have long been thought to stabilize plasmids by killing cells that fail to inherit a plasmid copy through a phenomenon called post-segregational killing (PSK) or addiction. While this model has been widely accepted, no direct observation of PSK was reported in the literature. Here, we devised a system that enables visualization of plasmid loss and PSK at the single-cell level using meganuclease-driven plasmid curing. Using the *ccd* system, we show that cells deprived of a *ccd*-encoding plasmid show hallmarks of DNA damage, *i*.*e*. filamentation and induction of the SOS response. Activation of *ccd*-triggered cell death in most plasmid-free segregants, although some intoxicated cells were able to resume growth, showing that PSK-induced damage can be repaired in a SOS-dependent manner. Damage induced by *ccd* activates resident lambdoid prophages, which potentiate the killing effect of *ccd*. The loss of a model plasmid containing TA systems encoding toxins presenting various molecular mechanisms induced different morphological changes, growth arrest and loss of viability. Our experimental setup enables further studies of TA-induced phenotypes and suggests that PSK is a general mechanism for plasmid stabilization by TA systems.

## Introduction

Plasmids are key drivers of genome evolution by promoting gene shuffling via horizontal gene transfer (1). In addition to cargo accessory genes that provide beneficial ecological traits under selective conditions, plasmids encode ‘core’ genes allowing for copy-number control, multimer resolution and partitioning in daughter-cells ensuring their stable maintenance in growing bacterial populations (2). Along with these maintenance functions, plasmids often encode toxin-antitoxin (TA) modules which are generally composed of two genes encoding a toxin and its cognate antitoxin (3–6). While these systems were originally discovered on the F and R1 plasmids in the 1980’s (7–9), the combination of comparative genomics, biochemistry and structural biology led to the identification and characterization of dozens of such systems encoded in bacterial plasmids and chromosomes (3–6). TA systems turned out to be widespread, employing toxins with highly diverse activities and antitoxins of different nature and mode of action (4, 10). The latter served as the basis for establishing TA systems classification and we currently distinguish eight different classes among which type I (RNA antitoxin inhibiting translation of its cognate toxin) and type II (protein antitoxin sequestering its cognate toxin) are the best-characterized (4).

The first type II TA module was identified on the *Escherichia coli* F plasmid. Works from several groups showed that this large conjugative plasmid encodes a two-gene locus (*ccd* for <u>c</u>ontrol of <u>c</u>ell <u>d</u>ivision) that prevents plasmid loss (8, 9). This type II TA system is composed of the CcdA antitoxin and CcdB toxin (9). An initial model proposed that upon decrease of F copy-number, inhibition of cell division by CcdB would allow plasmid replication and restauration of the appropriate number of segregational units to ensure successful plasmid partitioning in nascent daughter cells, therefore coupling plasmid replication to cell division (9). Population-level studies using a thermosensitive replicon carrying the *ccd* system revealed a plateauing in viable cell counts at non-permissive temperature, indicating that this locus could inhibit growth in plasmid-destabilizing conditions (9, 11). However, total cell counting under these conditions revealed a large population of non-viable cells concomitantly with filamentation, a phenotype reminiscent of DNA damage and induction of the SOS response (11, 12). Separation of filamentous cells by Percoll gradient showed that these cells are not viable, suggesting that *ccd* triggers DNA damage and cell death in plasmid-free segregants (11). This locus was ultimately named *ccd* for <u>c</u>ontrol of <u>c</u>ell <u>d</u>eath (13). Subsequent work established that the CcdB toxin poisons DNA gyrase, leading to the formation of double stranded DNA breaks (DSBs) and induction of the SOS response, supporting the filamentation phenotype (13, 14). Simultaneously to the discovery of the *ccd* system, another TA system was identified on the R1 plasmid (7). The type I *hok-sok* system was shown to comprise an antitoxin RNA (*sok*, <u>s</u>uppression <u>o</u>f host <u>k</u>illing) that inhibits translation of the toxin (*hok*, <u>ho</u>st <u>k</u>illing) (7). Using comparable replication thermosensitive replicons, the *hok-sok* locus was found to induce a plateau in viable cell counts at non-permissive temperature, in an analogous manner to the *ccd* locus (7). This was accompanied with a loss of cytosolic content, which indicated that this system triggers lysis of plasmid-free segregants through a molecular mechanism that differs from *ccd* (7). It was subsequently showed that the Hok toxin is small pore-forming protein that inserts into inner membrane and indeed provokes cell lysis (15). These pioneering studies laid the formulation of the post-segregational killing (PSK) model or addiction as quoted later on, in which TA systems favor plasmid retention in populations by inducing cell death in plasmid-free segregants (7, 11, 16). It was further showed that the molecular basis of addiction relies on different stability between the components, antitoxins being labile (17–19). While antitoxins and toxins are constantly replenished in plasmid-containing cells, loss of TA-encoding genes in plasmid-free daughter cells would lead to the depletion of the unstable antitoxin, liberation of the toxin and thus, target corruption and cell death (17–19).

While the addiction model was elaborated from fragmentary observations, it became an accepted paradigm regarding the mechanisms by which TA systems promote plasmid retention (3–6). However, models in which TA systems regulate plasmid replication or segregation continued to be proposed throughout the years. For example, overexpression of the Kid toxin from plasmid R1 was shown to uncouple DNA replication and cell division to facilitate plasmid inheritance, as initially proposed for the *ccd* system (20). Another study showed that the omega-epsilon-zeta tripartite system from *Streptococcus* plasmid pSM19035 was able to regulate plasmid copy number through the Omega gene product, which encodes the repressor component of this tri-partite TA system (21). Similarly, the PrpA antitoxin of the PrpAT system was shown to inhibit replication of the *Pseudoalteromonas* plasmid pMBL6842 by competing with the plasmid-encoded replication initiator for iteron sequences (22). However, in all those cases, conditions that alter regulator or antitoxin amounts are unclear, with the mechanism allowing these TA systems to control plasmid stability by mechanism other than PSK remain elusive. Since PSK is not mutually exclusive with such control mechanisms, it is not excluded that TA systems could stabilize plasmids by means other than PSK. Here we used time-lapse fluorescence microscopy to re-examine the PSK model by following the plasmid loss events in real time at the single-cell level. We first observed segregation of a fluorescent protein-tagged mini-F plasmid carrying the *ccd* locus. We provide direct evidence that this system does not prevent plasmid loss *per se*, but rather induces SOS response after plasmid loss, confirming that *ccd* is activated in plasmid-free segregants. To increase the loss frequency and facilitate our analysis, we engineered a unique system in which plasmid curing is forced through digestion by the I-SceI endonuclease. Using this system, we show that *ccd* triggers cell death in the majority of plasmid-free segregants, with some cells being able to escape killing due to SOS-dependent repair of CcdB-induced DNA damage. Curing plasmids encoding six other TA systems presenting various toxicity mechanisms also resulted in cell death, although these systems were much more lethal than *ccd*. Altogether, our work establishes an experimental system that enables to study TA activation at the single cell level and showcases the first live observation of PSK, establishing this phenomenon as a general mechanism that explains retention of TA-encoding replicons.

## Results

### Steady-state loss of mini-F plasmids reveals activation of the ccd toxin-antitoxin system in plasmid-free segregants

We first examined the unperturbed segregation of a fluorescently labeled mini-F replicon at the single-cell level. This replicon is actively segregated by the *sopABC* system, a type Ia partition system. SopB binds the *sopC* centromeric repeats to form partition complexes, which are segregated through the ATPase activity of SopA (23, 24). We fused SopB to mNeongreen, a bright monomeric fluorescent protein, allowing to follow partition complex localization with minimal perturbation of plasmid segregation as described previously (**Figure 1A**) (25, 26). Fluorescence was localized in foci along the medial axis of cells, confirming the functionality of our reporter (Figure 1A) (25, 26). At the population level, this plasmid was lost at a rate of 0.09% per generation, which is comparable to what was observed with other mini-F derivatives where SopB is tagged with a fluorescent protein (**Figure 1B**) (25, 26). Introduction of the *ccd* system in this plasmid lead to undetectable loss levels, confirming the plasmid-stabilizing property of this system (**Figure 1B**). To rule out any effect of *ccd* on segregational stability through an increase in plasmid copy number, the number of SopB-mNG foci per cell as a proxy for plasmid copy number was quantified. No difference was detected between an empty vector and a *ccd*-encoding plasmid (**Figure 1C**). DNA quantification by real-time polymerase chain reaction confirmed that *ccd* does not affect the copy number of our mini-F model plasmid (**Supplementary Figure 1**).

**Figure 1:**
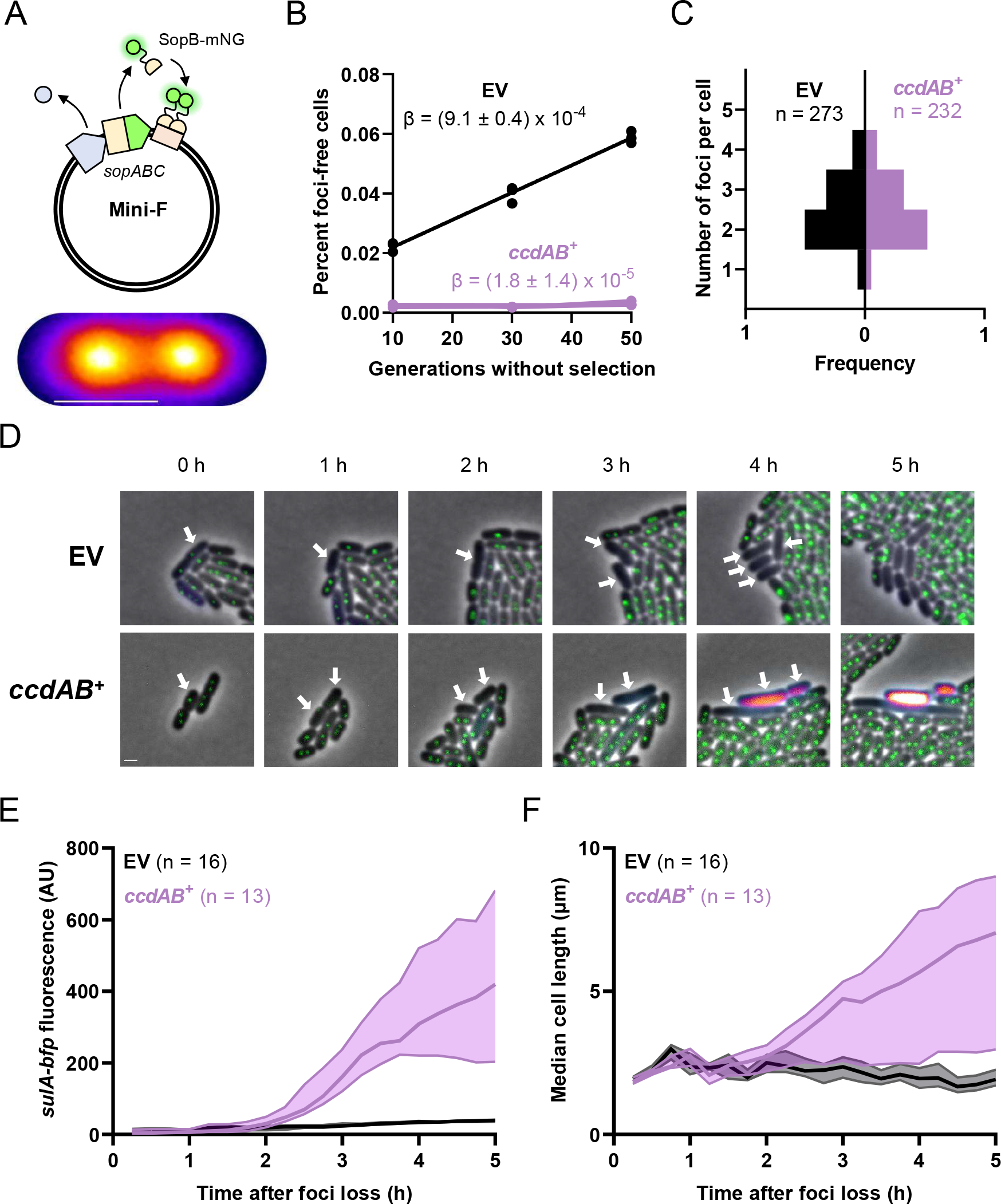
The *ccd sys*tem induces the SOS response in plasmid-free segregants. **A:** Illustration of the mini-F tracking system. Top: The SopB protein from the native partition system of F was fused to the mNeongreen fluorescent protein in the pNF03 mini-F plasmid. Binding of SopB-mNeongreen to the *sopC* centromere leads to the formation of green-fluorescent foci that enables positional tracking of the plasmid. Bottom: average SopB-mNeongreen fluorescence intensity of 115 cells displaying two SopB-mNeongreen foci. Scale bar is 1 μm. **B:** Loss mini-F plasmids in continuous culture. FN042 cells transformed with pNF03 (EV) and pNF03*ccd* (*ccdAB* ^+^) were grown to exponential phase in MOPS medium containing 0.4 % maltose and 15 μg/ml chloramphenicol to promote mini-F retention before being diluted 1000x in the same medium without antibiotic. Cells were diluted 1000x every 12 h and allowed to grow for 10 generations every cycle. The proportions of foci-free cells were counted at indicated timepoints and a linear regression was fit to the data of three independent replicates to obtain plasmid loss rate per generation (β). **C:** Copy number of mini-F vectors. FN042 cells transformed with pNF03 (EV) and pNF03*ccd* (*ccdAB* ^+^) grown on MOPS medium containing 0.4 % maltose were imaged on agarose pads. The numbers of SopB-mNeongreen foci were counted for the indicated number of cells. **D-F:** Time-lapse fluorescence microscopy analysis of unperturbed mini-F loss. FN042 cells transformed mini-F vectors pNF03 (EV) or pNF03*ccd* (*ccdAB* ^+^) imaged by time-lapse fluorescence microscopy every 15 min on agarose pads made with MOPS medium containing 0.4 % maltose. **D:** Representative micrographs of mini-F loss events. The SopB-mNeongreen fusion that localizes the plasmid is shown in green while the *sulA-bfp* fusion that reports the SOS response is shown with a fire LUT. Time 0 corresponds to the last timepoint a SopB-mNeongreen foci was visible in arrow-indicated cells. Scale bar is 1 μm. **E-F:** Quantification of median *sulA-bfp* fluorescence (**E**) and cell size (**F**) over time in the indicated numbers of plasmid free segregants after foci loss (time 0). Shaded areas represent interquartile ranges.

We imaged 3.2x10^4^ live divisions of cells carrying a *ccd*-deficient mini-F plasmid as well as 2.9x10^4^ divisions of cells carrying a *ccd*-encoding plasmid. Similar numbers of division events resulting in plasmid-free segregants were observed for the empty and *ccd*-encoding plasmids (29 and 31, respectively), suggesting that *ccd* does not affect the segregation of its replicon. A total of 16 *ccd*-deficient and 13 *ccd*-positive plasmid-free segregants were imaged in optimal conditions. We quantified cell length and induction of the SOS-response, using a *sulA-bfp* transcriptional reporter for the latter (**Figure 1D**) (27). A gradual increase in BFP fluorescence was observed after two hours in cells that lost a *ccd*-encoding plasmid but not in those carrying an empty vector, confirming that the SOS response is induced post-segregationally (**Figure 1E**). Filamentation, a hallmark of the SOS response (28), was observed in *ccd* plasmid-free segregants, synchronously with *sulA-bfp* induction (**Figure 1D**). Cells reached a median cell length of 7 μm 5 hours after plasmid loss, compared to 2 μm for cells that lost an empty plasmid (**Figure 1F**). Our results therefore demonstrate that *ccd* does not prevent plasmid mis-segregation but rather triggers DNA damage that leads to induction of the SOS response in plasmid-free segregants. Moreover, our single-cell data showcase the first live evidence of TA system activation in conditions where toxin concentrations are not ectopically manipulated.

### Meganuclease-mediated curing of mini-F plasmids induces ccd-mediated killing

Because steady-state loss of the mini-F plasmid is a rare occurrence, plasmid-free segregants quickly get outcompeted and drowned in plasmid-bearing siblings, rendering tracking of their fate difficult. Indeed, while the aforementioned setup enabled us to visualize *ccd* activation and induction of the SOS response after plasmid loss, it failed to provide information regarding the viability of these plasmid-free segregants. To circumvent these limitations, we devised a system to induce synchronous plasmid curing in the whole population. We introduced a I-SceI cutting site on a mini-F plasmid and provided an arabinose-inducible *SCE1* gene *in trans* (**Figure 2A**). In this setup, addition of arabinose induces production of I-SceI, which specifically recognizes its cognate cutting site on the plasmid, leading to plasmid cleavage and degradation by endogenous exonucleases such as RecBCD (29). The absence of Chi sites in this plasmid should also prevent loading of RecA by RecBCD and therefore limit plasmid recombinational repair and induction of the SOS response (29). Quantitative polymerase chain reaction showed a 99.8 % decrease of plasmid DNA levels 3 hours after arabinose induction of SCE1, confirming the functionality of our plasmid curing system (**Supplementary Figure 2**). Curing an empty plasmid did not affect viability since plating cells on medium containing arabinose did not decrease colony-forming units (CFU) compared to medium containing glucose and kanamycin, showing that our plasmid curing system has a minimal impact on cell growth and viability. (**Figure 2B**). On the other hand, curing a *ccd*-encoding plasmid led to a 68% decrease in cell viability, showing that loss of a *ccd*-encoding plasmid induces cell death in most of the population (**Figure 2B**). Abolishing the toxicity of CcdB by substituting its 100^th^ residue from glycine to glutamate (14, 30) or by using a *gyrA462* mutant, which is not intoxicated by CcdB (13, 14), restored viability and confirmed that gyrase poisoning is required for *ccd*-induced post-segregational killing (**Figure 2B**). Likewise, *ccd*-induced killing after curing is abolished in a *lon* mutant, confirming a requirement of CcdA degradation by Lon to enable PSK as previously suggested (**Figure 2B**) (17).

**Figure 2:**
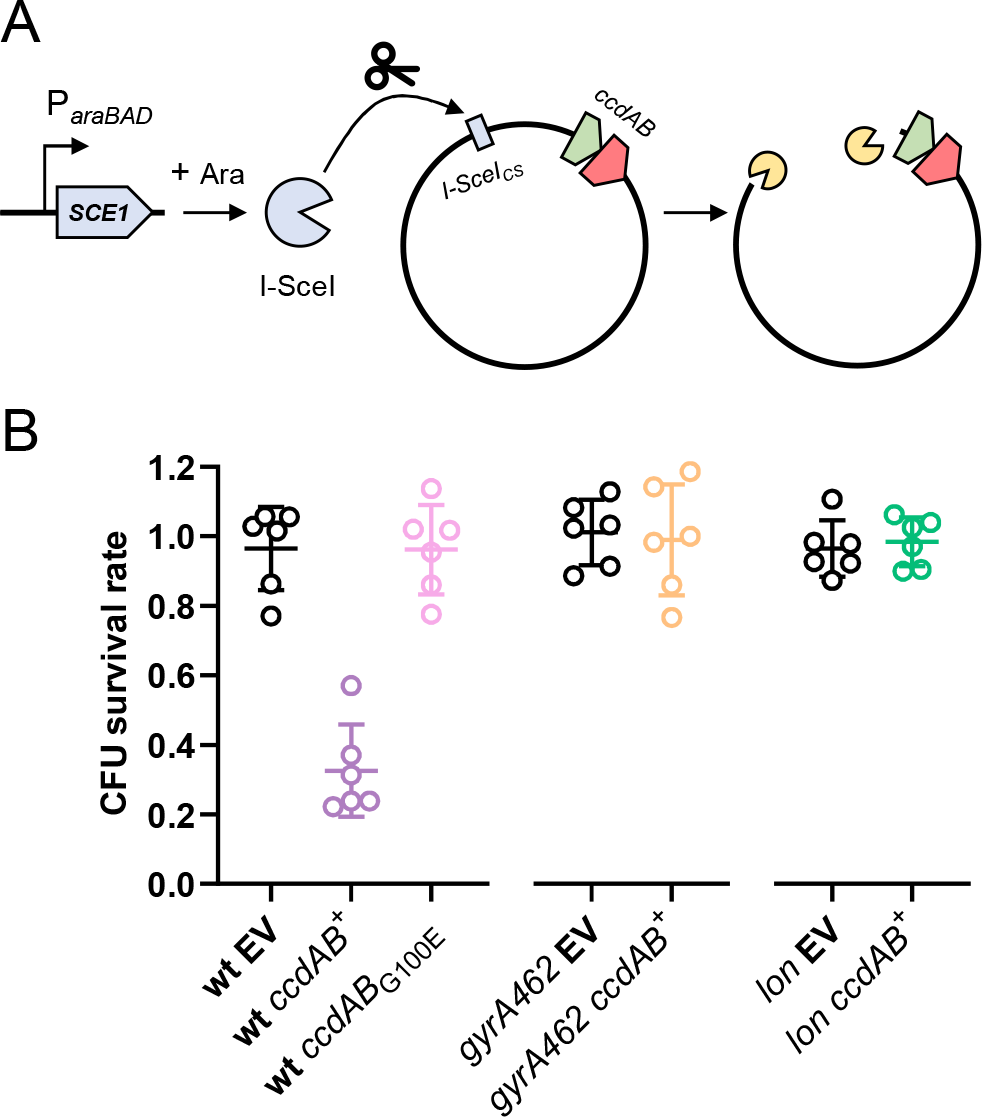
I-SceI-mediated curing reveals post-segregational killing. **A:** Schematic representation of the plasmid curing system. The SCE1 gene is under the control of the *araBAD* arabinose-inducible promoter on a low-copy plasmid (pSce), which allows production of the I-SceI restriction enzyme when cells are grown in the presence of arabinose. Plasmid pNF04, which is used to clone TA systems, contains a I-SceI cutting site (*I-SceI*_*CS*_), which allows its digestion and curing by the restriction enzyme when arabinose is added to the medium. B: Viability loss by ccd under plasmid curing conditions. Cells transformed with pSce and pNF04 derivates (pNF04, EV; pNF04ccd, *ccdAB*^+^ ; pNF04*ccdGE*; *ccdAB*_*G100E*_) were grown to exponential phase in MOPS medium containing 0.4 % glucose, 25 μg/ml kanamycin and 20 μg/ml chloramphenicol, serially diluted and spotted on M9 plates containing 20 μg/ml chloramphenicol and either 0.4% glucose and 25 μg/ml kanamycin to promote plasmid retention, or 0.1% glucose and 0.3% arabinose to promote plasmid curing. Data represents the mean and standard deviation of three independent experiments.

Altogether, by specifically triggering plasmid curing we provide direct evidence that *ccd* is activated in plasmid deprived cells, eliminating most of this population and thereby increasing plasmid retention at the population level. Moreover, we validate that plasmid curing and TA activation can be induced *à la carte* by I-Sce-mediated cleavage, therefore rendering this experimental setup highly suitable and attractive to study TA activation at physiological concentrations and identify factors regulating this process.

### Live imaging of plasmid curing reveals ccd-mediated post-segregational killing

To further investigate the phenotypical changes brought by *ccd* upon plasmid curing, the plasmid curing system used above was adapted for use in time-lapse microscopy analysis. By growing cells with arabinose to induce I-SceI production, a progressive loss of SopB-mNeongreen foci produced by our model mini-F plasmid can be observed (**Figure 3A, Supplementary Figure 3**). We imaged the growth of 122 microcolonies during SCE1 induction, of which 67% had lost the SopB foci after 1 hour. After 3 hours, 100 % of those microcolonies had lost their foci, confirming that I-SceI production induces plasmid loss at the single cell level (**Supplementary Figure 3, Supplementary Movie 1**).

**Figure 3:**
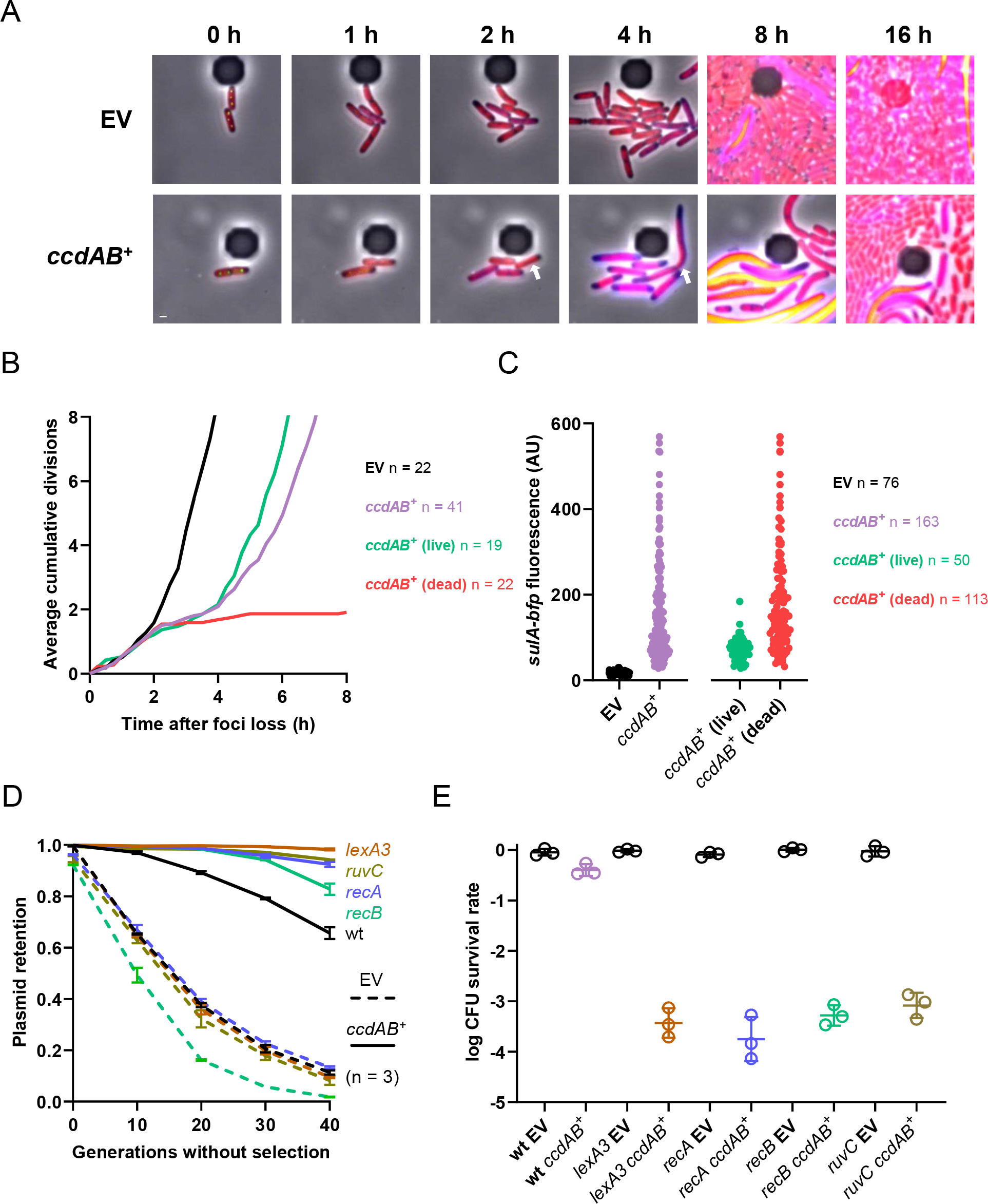
Live imaging of *ccd*-mediated post-segregational killing. **A-C:** Time-lapse fluorescence microscopy analysis of I-SceI-mediated plasmid curing. FN053 cells were transformed with pSce and either pNF04 (EV, top) or pNF04*ccd* (bottom) plasmids, grown on MOPS medium containing 0.4 % glucose, 25 μg/ml kanamycin and 20 μg/ml chloramphenicol, then loaded on microfluidic chips, perfused with MOPS medium containing 0.4 % arabinose and 20 μg/ml chloramphenicol to induce plasmid curing and imaged by time-lapse fluorescence microscopy every 15 min. **A:** Representative micrographs of plasmid-free segregants following I-SceI-mediated curing. SopB-mNeongreen is shown in green, the HU-mCherry fusion that localizes the chromosome is shown in red and the *sulA-bfp* is shown in fire LUT. A cell surviving the effects of *ccd* is shown by a white arrow. Scale bar is 1 μm. **B:** Quantification of divisions by microscopy after plasmid loss. The number of cumulative divisions for observed plasmid loss events were quantified at each 15 min timepoint after SopB foci loss and averaged over the indicated number of imaged plasmid loss event (n). Fate of daughter cells from plasmid curing events was classified as dead (failure to form microcolonies) or survivors (formation of microcolones). **C:** Quantification of *sulA-bfp* median fluorescence intensity in daughter cells of plasmid-free segregants 4 hours after SopB foci loss. **D:** Plasmid-retention in DSB-repair-deficient mutants. Indicated strains transformed with pNF06 (dashed lines) or pNF06*ccd* (solid lines) were continuously grown in MOPS medium containing 0.4 % glucose and sampled at indicated times during exponential phase to measure the proportion of fluorescent cells by flow cytometry. Data represents the mean and standard deviation of three independent experiments. **E:** Viability of DSB-repair-deficient mutants under plasmid curing conditions. Indicated mutants transformed with pSce and either pNF04 (black) or pNF04*ccd* (colors) were grown to exponential phase in MOPS medium containing 0.4 % glucose, 25 μg/ml kanamycin and 20 μg/ml chloramphenicol, serially diluted and spotted on M9 plates containing 20 μg/ml chloramphenicol and either 0.4% glucose and 25 μg/ml kanamycin to promote plasmid retention, or 0.1% glucose and 0.3% arabinose to promote plasmid curing. Data represents the geometric mean and standard deviation of six independent experiments.

As a proxy for CcdB activity, we quantified cumulative divisions and induction of the *sulA-bfp* reporter in plasmid-free segregants after plasmid curing. Curing an empty plasmid did not affect cell growth, which continued to divide exponentially after arabinose addition (**Figure 3A, B, Supplementary Movie 2**). After losing a *ccd*-encoding plasmid, cells kept on dividing normally for 2 hours (**Figure 3A, B**). After this timepoint, division rate plateaued and *sulA* induction was detected in *ccd* plasmid-free segregants but not in cells that lost the control plasmid (**Figure 3A-C**). However, division rate accelerated after the third hour of induction, which reflected a resumption of division and growth in part of the population (**Figure 3B**). Out of 41 analyzed loss events of a *ccd*-encoding plasmid, 19 (46 %) resulted in division resumption and formation of a microcolony post plasmid-loss, underlying the survival of these plasmid-free segregants (**Figure 3B**). On the other hand, 22 (54 %) of these plasmid-free segregants failed to resume division and to produce microcolonies (**Figure 3B**). Both populations had detectable blue fluorescence produced by the *sulA-bfp* reporter four hours after plasmid loss, suggesting that CcdB induces DNA damage in both subpopulations (**Figure 3C**). The absence of *sulA-bfp* induction at this timepoint in cells that lost an empty plasmid (17 ± 5 AU) also confirmed that cleavage of our model plasmid did not induce the SOS response (**Figure 3C**). While *sulA-bfp* induction levels showed a broad distribution in the bulk of the population (151 ± 115 AU), surviving cells were consistently located at the lower end of this spectrum (73 ± 28 AU). A lower induction of the SOS response could indicate less DNA damage endured by surviving cells and repair of CcdB-induced DSBs (**Figure 3C**).

We then assessed whether the DSB repair machinery was involved in survival to *ccd*-mediated post-segregational killing. DSB repair first requires end resection by the RecBCD nuclease, loading of RecA on single-stranded DNA (ssDNA) and activation of the SOS response by RecA-mediated autocleavage of the LexA repressor (28, 29, 31). RecA-ssDNA nucleoprotein filaments mediate pairing of the break with a sister locus. Subsequent strand invasion and homology-mediated repair of the break generate Holliday junctions that need to be resolved by the RuvABC complex (32). We first measured the stability of a partition-deficient plasmid in mutants deficient for every step of this repair process, which include a non-inducible *lexA3* mutant as well as *recA, recB* and *ruvC* mutants. After 40 generations of culture without selection for plasmid retention, only 11% of the cells retained fluorescence (**Figure 3D**). A *ccd*-encoding plasmid showed better yet limited retention, with 66% of the cells remaining fluorescent after 40 generations of culture (**Figure 3D**). The empty vector was lost at similar rates in *lexA3, recA* and *ruvC* mutant while loss was slightly accentuated in a *recB* mutant (**Figure 3D**). However, a *ccd*-encoding plasmid displayed better retention in all these mutants when compared to a wild-type strain, with all mutants, *recB* excepted, showing a plasmid retention higher than 90% (**Figure 3D**), suggesting that survival from *ccd*-mediated PSK requires repair of DSBs. Consistently with these results, I-SceI-mediated curing of a *ccd*-encoding plasmid reduced viability by 3 to 4 orders of magnitude in *recA, recB, ruvC* and *lexA3* compared to a wild-type strain, while curing of an empty plasmid did not affect viability in these mutants (**Figure 3E**). We therefore show that *ccd* is activated in all plasmid-free segregants, in which it induces DNA damage. However, this activation only results in a partial elimination of cured cells. Mutants inactivated for the repair of DSBs are more efficiently killed by *ccd* by several orders of magnitude, suggesting that survival to *ccd*-mediated PSK is facilitated by the repair of CcdB-induced DNA lesions. Moreover, these data validate the functionality of our experimental system to study TA activation and subsequent phenotypes at the single-cell level.

### Cooperativity between ccd and lambdoid prophages

To further study *ccd*-induced phenotypes and illustrate the suitability of our experimental setup, we characterized the activation of *ccd* and its downstream consequences in backgrounds that carry lysogen lambdoid prophages. Since DNA double-stranded breaks are known to derepress lambdoid phages and induce the lytic production of viral particles (31), we investigated whether *ccd*-induced PSK in lambda lysogens could induce lysis and whether this lysis could, in turn, promote retention of a *ccd*-encoding plasmid in lysogens. While *ccd*-mediated PSK in a non-lysogen background triggered lysis in 0.6% of the population (**Supplementary Movie 3**), imaging of plasmid curing in lambda lysogens reveals a post-segregational induction of lysis in most cells that lost a *ccd*-encoding plasmid (58.0 %) but not in those that lost an empty vector (0.8%) (**Figure 4A-B, Supplementary Movie 3-4**). Production of viral particles was quantified to confirm induction of the lytic cycle in *ccd* plasmid-free segregants. Lambda lysogens grown on MOPS glucose medium produced 10^3^-10^4^ plaque-forming units (PFU) whether they were transformed with an empty or *ccd*-encoding plasmid (**Figure 4C**). Growth on arabinose to induce plasmid curing increased PFU yields by three orders of magnitude for cells that cured a *ccd*-encoding plasmid, but not for those that cured an empty plasmid, showing that loss of a *ccd*-encoding plasmid triggers entry into the lytic cycle followed by virion production (**Figure 4C**).

**Figure 4:**
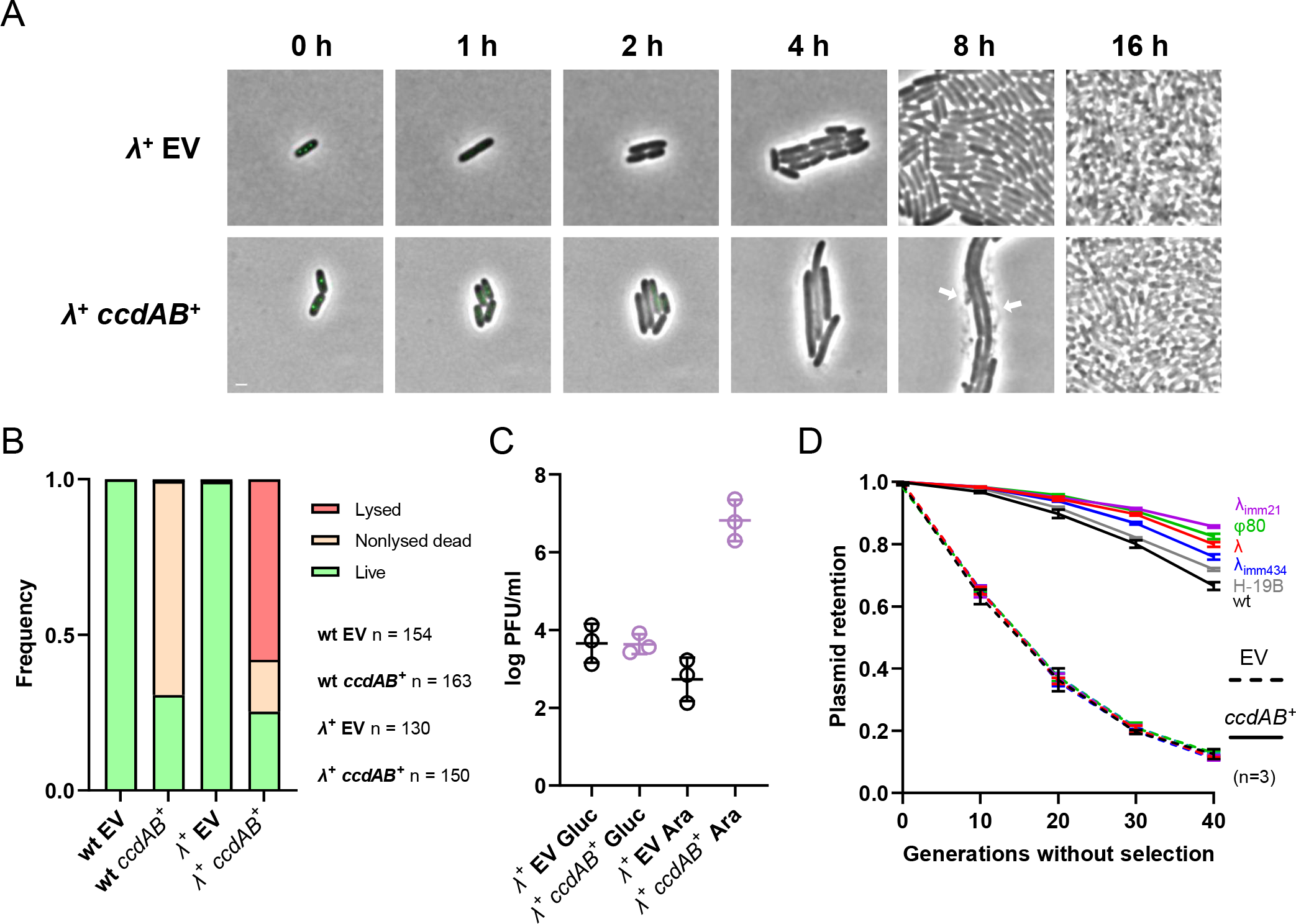
Cooperativity between *ccd and* lambdoid prophages. **A-B:** Time-lapse fluorescence microscopy analysis of I-SceI-mediated plasmid curing in lambda lysogens. MG1655 cells were lysogenized with lambda and transformed with pSce and either pNF04 (EV, top) or pNF04*ccd* (bottom) plasmids, grown on MOPS medium containing 0.4 % glucose, 25 μg/ml kanamycin and 20 μg/ml chloramphenicol, spotted on agarose pads with MOPS medium containing 0.4 % arabinose and 20μg/ml chloramphenicol to induce plasmid curing and imaged by time-lapse fluorescence microscopy every 15 min. **A:** Representative micrographs of plasmid-free segregants following I-SceI-mediated curing. SopB-mNeongreen is shown in green. Scale bar is 1 μm. **B:** Quantification of survival and lysis in plasmid-free segregants in non-lysogens cells grown in microfluidics (as in Figure 3A) or lambda lysogens grown on agarose pads as described above. Fate of daughter cells from plasmid curing events was classified as lysed (disappearance or loss of phase contrast), nonlysed dead (failure to form microcolonies) or live (formation of microcolones) in the indicated number of cells (n). **C:** Production of viral particles by *ccd-*induced PSK. Lysogens transformed with pSce and either pNF04 (EV) or pNF04*ccd* were grown to exponential phase in MOPS medium containing 0.4 % glucose, 25 μg/ml kanamycin and 20 μg/ml chloramphenicol, and diluted 100x MOPS medium containing 20 μg/ml chloramphenicol and either glucose (Gluc) or arabinose (Ara) as sole carbon source. Plaque-forming units (PFU) were estimated from lysates after 16 hours of growth. **D:** Plasmid-retention in lysogens. Lysogens of indicated phages transformed with pNF06 (dashed lines) or pNF06*ccd* (solid lines) were continuously grown in MOPS medium containing 0.4 % glucose and sampled during exponential phase at indicated times to measure the proportion of fluorescent cells by flow cytometry. Data represents the mean and standard deviation of three independent experiments.

We next investigated whether lysogeny could enhance *ccd*-mediated PSK by inducing lysis in cells that would have otherwise survived intoxication by CcdB following plasmid loss. Our microscopy assays could detect a marginal reduction of survival to *ccd*-mediated PSK in lambda lysogens (25.3%) compared to non-lysogens (30.7%) (**Figure 4B**). However, plating on arabinose versus glucose medium failed to detect such a small difference, likely due to the intrinsic error of ten-fold dilutions used in this assay, which is more suited to detect differences on a logarithmic scale (**Supplementary Figure 4**). To robustly evaluate the effect of lysogeny on PSK and plasmid stabilization by *ccd*, we quantified the retention of a partition-defective mini-F plasmid in lambda lysogens by flow cytometry over forty generations. Plasmid retention was also investigated in other lambdoid lysogens (ϕ80 and H-19B) as well as in heteroimmune lysogens (λ_imm434_ and λ_imm21_). All lysogens showed a similar retention of the empty plasmid as a wild-type strain (**Figure 4D**). On the other hand, retention of a *ccd*-encoding plasmid was improved with variable efficiencies in lysogens (**Figure 4D**). The λ_imm21_ lysogen showed the greatest retention with 86% fluorescent cells after four days of culture while a H19-B lysogen had the poorest retention with 72% fluorescent cells, compared to a non-lysogen that displayed only 66% fluorescent cells (**Figure 4D**). Lysogeny by lambdoid prophages therefore increased PSK by *ccd*, likely through the induction of a lytic cycle in plasmid-free segregants that would have otherwise repaired CcdB-induced DNA damage.

### Post-segregational killing is a conserved mechanism for TA-mediated plasmid stabilization

Although *ccd* and *hok-sok* were the first identified TA systems, other plasmid-encoded TA with various toxic activities were subsequently identified. These include the *parDE* system from IncP-1 plasmids RK2 and RP4, in which the toxin is a gyrase inhibitor structurally unrelated to CcdB (33, 34) ; the *higBA* system from the Rts1 plasmid, in which the toxin cleaves messenger RNAs in a translation-dependent manner by entering the ribosomal A site (35, 36), the *vapBC* system from the plasmid pINV of *Shigella flexneri*, in which the toxin cleaves the anticodon loop of the fMet-tRNA (37–39); and the *phd-doc* system from phage P1, in which the toxin inactivates translation elongation factor Tu by phosphorylation (16, 40) and. (**Figure 5A**). So far, killing by these systems has only been studied using toxin overproduction, with the assumption that activation of toxins from their native loci during PSK would trigger comparable toxicity and killing (3, 33, 35, 37, 40). As with *ccd*, these systems inhibited growth at non-permissive temperatures when cloned in temperature-sensitive plasmids, suggesting that these systems are activated when plasmid replication is compromised (16, 34, 36, 39).

**Figure 5:**
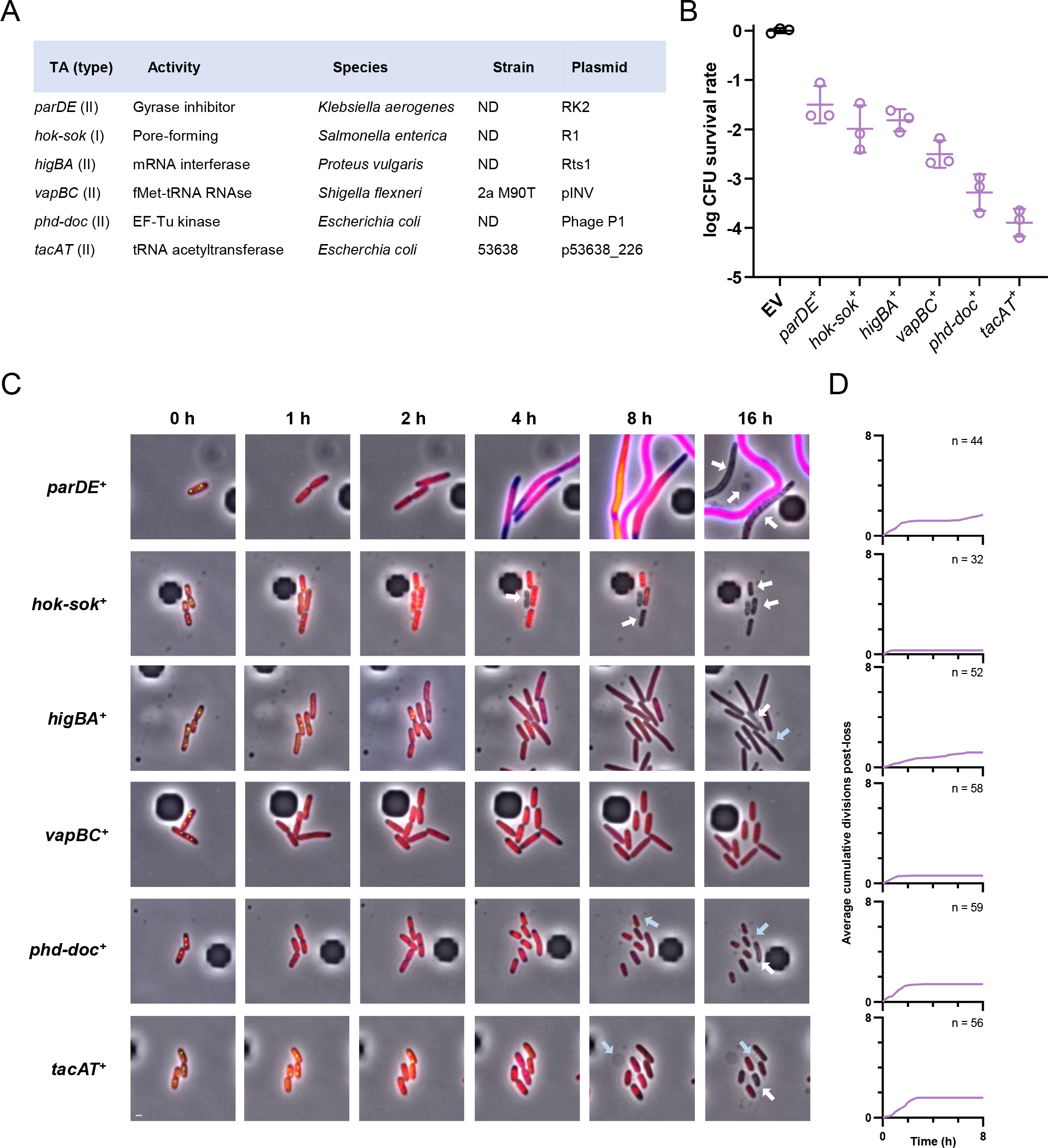
Conservation of post-segregational killing across TA families. **A:** Recapitulative table of tested TA systems and their origin. ND : not determined. B: TA-induced viability loss under plasmid curing conditions. Cells transformed with pSce and pNF04 derivatives containing the indicated TA systems were grown to exponential phase in MOPS medium containing 0.4 % glucose, 25 μg/ml kanamycin and 20 μg/ml chloramphenicol, serially diluted and spotted on M9 plates containing 20 μg/ml chloramphenicol and either 0.4% glucose and 25 μg/ml kanamycin to promote plasmid retention, or 0.1% glucose and 0.3% arabinose to promote plasmid curing. Data represents the geometric mean and standard deviation of three independent experiments. **C-D:** Time-lapse fluorescence microscopy analysis of I-SceI-mediated plasmid curing. FN053 cells were transformed with pSce and either pNF04 (EV, top) or pNF04*ccd* (bottom) plasmids, grown on MOPS medium containing 0.4 % glucose, 25 μg/ml kanamycin and 20 μg/ml chloramphenicol, then loaded on microfluidic chips perfused with MOPS medium containing 0.4 % arabinose and 20 μg/ml chloramphenicol at time 0 to induce plasmid curing. Cells were imaged by time-lapse fluorescence microscopy every 15 min. **C:** Representative micrographs of plasmid-free segregants following I-SceI-mediated curing. SopB-mNeongreen is shown in green, the HU-mCherry fusion that localizes the chromosome is shown in red and the *sulA-bfp* is shown in fire LUT. Scale bar is 1 μm. White arrows show loss of cytosolic content while blue arrows show blebbing. **D**: Quantification of divisions by microscopy after plasmid loss. The number of cumulative divisions for observed plasmid loss events were quantified at each 15-minute timepoint after SopB foci loss and averaged over the indicated number of imaged plasmid loss event (n).

We therefore assessed whether these systems could trigger PSK when produced through their native loci by cloning them in our curable mini-F vector. We also cloned a plasmid-encoded *tacAT* system that showed 90% identity at the protein level with the well described *tacAT* system from Salmonella enterica, which acetylates Gly-tRNA^Gly^ (41) (**Figure 5A**).

Curing plasmids encoding each of these six TAs reduced viability by several orders of magnitude, suggesting that these TAs kill plasmid-free segregants more efficiently compared to *ccd* (**Figure 5B**). Interestingly, these systems showed varying efficiencies of killing, with *vapBC* (3.6x10^−3^), *phd-doc* (6.4x10^−4^) and *tacAT* (1.4x10^−4^) displaying the lowest survival rates (**Figure 5B**). On the contrary, parDE (4.2x10^−2^), *hok-sok* (1.5x10^−2^) and *higBA* (1.7x10^−2^) showed lower killing efficiencies after curing, suggesting that these systems are less efficient at killing plasmid-free segregants (**Figure 5B**).

Microscopy analysis of plasmid-cured cells revealed that plasmid loss is correlated with division inhibition for all tested TA systems (**Figure 5C-D, Supplementary Movies 5-10**). The *parDE* system, which poisons gyrase, induced similar phenotypes as the *ccd* system, including induction of the SOS response and filamentation (**Figure 5C, Supplementary Movie 5**) . The *hok-sok* system induced rapid growth arrest followed by congregation of SopB-mNeongreen fluorescence on the edges of the cells ; loss of cytoplasmic content could be detected over the course of several hours, consistent with the previous observation of Hok-induced “ghost” cells (7) (**Figure 5C, Supplementary Figure 5, Supplementary Movie 6**). Translation-inhibiting TAs (*vapBC, phd-doc, higBA* and *tacAT*) induced growth arrest, although *higBA* plasmid-free segregants kept dividing at a slow pace throughout the experiment (**Figure 5C, Supplementary Movies 7-10**). Surprisingly, blebbing-like protrusions visible by phase contrast could be detected in *phd-doc, higBA* and *tacAT* plasmid-free segregants, suggesting that the toxins of this systems could perturbate envelope homeostasis by an unknown mechanism (**Figure 5C, Supplementary Figure 5, Supplementary Movies 7, 9 & 10**). Altogether, these results show that activation of various families of type I and II TA systems from their native loci result in cell death, demonstration that PSK is a conserved mechanism by which TA systems mediate plasmid addiction.

## Discussion

In this work, we designed a system that enables curing of plasmids as a mean to study PSK by toxin-antitoxin systems. Visual clues PSK were obtained in previous studies by imaging cultures several hours after destabilizing TA-encoding replicons, with *ccd* inducing filamentation and production of anucleate cells, and *hok-sok* inducing cell lysis (7, 11). Likewise, destabilization of Chromosome II (ChrII) in *Vibrio cholerae* through the deletion of its *parABS* system resulted toxicity and DNA damage, which were dependent on three ChrII-encoded *parDE* systems (42). However, live imaging of this phenomenon has never been reported. Here, we report the first live observation of this phenomenon on its full scale and in a systematic manner.

Our work first imaged thousands of cell divisions and plasmid partition events by microscopy, showing that the *ccd* system does not increase segregational stability of its mini-F replicon. Rather, loss of a plasmid that carried the *ccd* system induces the SOS response and cell filamentation, supporting earlier observations that *ccd* leads to DNA damage in plasmid-free segregants. To facilitate the study of such TA-induced phenotypes, we were able to induce quasi-synchronous curing of the mini-F plasmid in the whole population through I-SceI-mediated cleavage of the plasmid, allowing to visualize intoxication of plasmid-free segregants by CcdB. A significant portion of plasmid-free segregants were able to survive PSK by repairing CcdB-mediated double-stranded DNA breaks, which translated into a poor stabilizing capacity for this system. Accordingly, previous reports showed that *ccd* is a poor plasmid stabilizer compared to other TA systems like *vapBC, parDE* or *hok-sok* which promoted plasmid retention orders of magnitude higher than *ccd* (38, 43). Redundancy of *ccd* with other TA systems encoded on the same plasmid, i.e., *flm* (which is quasi-identical to *hok-sok*) and *srn* on F, as well as *vapBC* and *gmvAT* on pINV (38, 44), could lead to a partial decay in the activity of the *ccd* system, through mutations that reduce *ccd* expression or diminish CcdB binding to DNA gyrase. Plasmid hosts could also have acquired mutations in the *gyrA* gene that reduce CcdB toxicity to facilitate plasmid curing and/or reduce lethality induced by *ccd*. We also showed a potential interplay between *ccd* and lambdoid prophages where PSK mediated by *ccd* induces lysis and virion production, which enhances retention of a *ccd*-encoding plasmid.

By using I-SceI-mediated curing of mini-F vectors encoding several other TA systems, our work demonstrates that PSK is a conserved mechanism by which TA of type I and II systems stabilize their replicon. Time-lapse microscopy analysis allowed us to track single plasmid curing events, which lead to defects consistent with the activities of each tested system, whether it be topoisomerase-poisoning, pore-forming, or translation-inhibiting. While inhibiting translation using antibiotics (*e*.*g*. tetracyclines, aminoglycosides, macrolides and phenicols) is widely regarded as bacteriostatic (45, 46), translation-inhibiting TAs induced killing more efficiently compared to TAs whose activities can be regarded as bactericidal, such as gyrase poisoning or inner membrane permeation. Interestingly, translation-inhibiting TAs induce cell lysis (*higBA*) or envelope abnormalities (*phd-doc, tacAT*), suggesting that corruption of the translation machinery by these systems has secondary effects that affect envelope homeostasis, which are ultimately bactericidal. However, the molecular mechanisms underlying these secondary effects remain to be elucidated.

While the effects of toxin on bacterial physiology is studied using multicopy expression vectors (*e*.*g*. pBAD arabinose-inducible plasmids), which enable the ectopic production of biologically-irrelevant levels of toxin (3), the experimental setup we describe in this study allows to study intoxication by TA systems and its downstream effects by triggering these system through their canonical activation pathway, *i*.*e*. through the loss of TA-encoding genes. We present here that our setup can be used to study TA activation at the single cell level to assess morphological parameters as well as the activation of responses through the use of fluorescent reporters. Owing to the fact that I-SceI induces plasmid curing in the whole population, our setup can also be used to study phenotypical parameters at the population level, with global approaches like RNAseq, ribosome profiling or RNA end mapping being well suited to study the effect of translation-inhibiting toxins at biologically relevant toxin levels. Other biological parameters such as accumulation of reactive oxygen species, ATP levels as well as membrane integrity and polarization can be assessed to study the downstream effects of these toxins that ultimately result in cell death.

## Acknowledgements

We are grateful to Jean-Yves Bouet, Fernando de la Cruz and Christian Lesterlin for donating strains and plasmids. Work in the Van Melderen lab is supported by the International Brachet Stiftung, the Université Libre de Bruxelles (Actions Blanches), the FNRS-FRS (RICOTTAS, T.0209.22 PDR), the Wallonia Region (ALGOTECH, 1510598) and the ‘Actions de Recherche Concertées’ (ARC, 2018-2023).

## Materials & Methods

### Strain & plasmid constructions

Plasmids used in this study are detailed in **Supplementary Table 1**. Plasmids were constructed by standard restriction ligation methods using T4 DNA ligase (NEB) or using the NEBBuilder assembly kit (NEB). PCR reactions were performed using Q5 DNA polymerase (NEB) or PrimeSTAR MAX (Takara). Oligonucleotides primers used in this study are detailed in **Supplementary Table 2**.

The trackable mini-F vector that yielded pNF03 was constructed by ligating a synthetic mNeongreen-encoding gene (47) at the SacI & PacI sites of pJYB240 (48). pNF03*ccd* was constructed by restoring a frameshift in *ccd*B using primers ApaLI-ccdB F & ApaLI-ccdB R then by digesting the PCR product with ApaLI and recircularizing it. pNF03 was constructed by deleting the *ccd*AB operon using primers delccd F and delccd R.

The pNF04 plasmid, a trackable mini-F vector that can be cleaved by I-SceI, was constructed by amplifying a mini-F replicon with a mNeongreen-tagged SopB from pNF03 using primers NotI-pNF04bb F and HindIII-pNF04bb R while a kanamycin resistance cassette was amplified from pUA66 (49) using primers HindIII-KmR F & NotI-KmR R. These two fragments were digested by NotI & ApaLI and ligated. A I-SceI cutting site was inserted into this vector using primers 04Sce F & 04Sce R, yielding pNF04.

The pSce plasmid, which allowed arabinose-induced production of I-SceI, was constructed using the NEBbuilder assembly by assembling a fragment amplified from pDL2655 (29, 50) using primers CmR-Sce F and CmR-Sce R and a pSC101 origin of replication amplified from pUA66 (49) using primers ori-Sce F and ori-Sce R, yielding pSce.

TAs were cloned in the AatII and HindIII sites of pNF04 and pNF05 using the following primers on their respective templates: AatII-*ccd* F & HindIII-*ccd* R for *ccd*AB from pNF03*ccd*, AatII-vap F & HindIII-vap R for *vapBC* from S. *flexneri* M90T genomic DNA, AatII-par F & HindIII-par R for *parDE* from RH8000, AatII-doc F & HindIII-doc R for *phd-doc* from phage P1vir, AatII-hok F & HindIII-hok R for *hok-sok* from BW27873 R1^+^, AatII-hig F & HindIII-hig R for *higBA* from a synthetic gene derived from pRts1, and AatII-tac F & HindIII-tac R for *tacAT* from a synthetic gene derived from *E. coli* 53638.

All strains used were isogenic to the MG1655 clone used as wild-type strain, in which relevant alleles were transduced. FRT-flanked resistance cassettes were excised using *flp* expression from pCP20 when applicable (see **Supplementary Table 3**) (51).

### Time-lapse microscopy analysis

Cultures for microscopy were prepared by diluting overnight cultures to OD_600nm_ 0.05 in MOPS medium (52) with indicated supplements. After reaching OD_600nm_ 0.5, these cultures were diluted 100x in the appropriate medium and either spotted on a sealed agarose pad (MOPS medium, 2% agarose, **Figure 1D-F & 4A**) or loaded in a CellASIC Onix plate (Merck) and perfused with MOPS medium containing 0.4% arabinose and 20 μg/ml chloramphenicol at 34.5 psi (**Figure 3A & 5C**). Microscopy experiments were performed using an Axio Observer Z1 microscope (Zeiss) equipped with a heating chamber, a motorized stage, a LED illumination system (Colibri 7, Zeiss) and a sCMOS camera (ORCA-Flash4.0 V2, Hamamatsu). mTagBFP2 was imaged using a 430 nm LED (12% intensity, 500 ms exposure), a 405/40 nm excitation filter and a 455/50 nm emission filter (Chroma). mNeongreen was imaged using a low power 511 nm LED (50% intensity, 2000 ms exposure), a 460/40 nm excitation filter and a 535/50 emission filter (Chroma). mCherry was imaged using a 590 nm LED (50% intensity, 500 ms exposure), a 530-585 excitation filter and a 615 nm long pass emission filter (Filterset 00, Zeiss). Images were taken every 15 min. Cells were outlined and segmented using MicrobeJ (53), with median intensity of fluorescence channels used as measurements of fluorescence after removal of the camera offset (100 bits).

### Plasmid retention assay by flow cytometry

Overnight cultures grown in MOPS medium with kanamycin were diluted to OD_600nm_ 0.05 in MOPS medium with 25μg/ml kanamycin and grown to exponential phase (OD_600nm_ 0.5). These cultures were then diluted 1000 × in MOPS medium without kanamycin and grown for indicated times before being processed by an Attune Nxt Flow cytometer. Green fluorescence was measured using a 488 nm laser and a 522/31 emission filter (Omega Filters). Cells were gated empirically to remove background signal and cell doublets were filtered out based on their higher side-scattering pulse area-to-height ratio.

### Post-segregational killing assay on plates

Overnight cultures were diluted to OD_600nm_ 0.05 in MOPS medium supplemented with 0.4% glucose, 20 μg/ml chloramphenicol, and 25 μg/ml kanamycin and grown at 37 °C. At OD_600nm_ 0.5, cells were serially diluted in PBS and plated on M9 plates (22 mM KH2PO4, 48 mM Na2HPO4, 18.8 mM NH4Cl, 8.6 mM NaCl, 2 mM MgSO4) supplemented either with 0.1% glucose and 0.3% arabinose to induce plasmid curing or 0.4% glucose and 25 μg/ml kanamycin to promote plasmid retention. These plates were then incubated for 30 h at 37 °C before colony counting.

### Real-time quantitative polymerase chain reaction

Samples of 1 ml were taken from cultures grown to OD 0.5 and centrifuged (5000 G, 5 min). After washing with 0,9 % NaCl, cell pellets were resuspended in 100 μl lysis buffer (10 mM Tris-HCl pH 8.0, 1 mM EDTA, 1 % Triton X-100, 0.5 % Tween 20) and boiled on a heating block for 10 min. Debris were pelleted (10000 RCF, 2 min) and supernatants were conserved at -20°C until further use. PCR mixes were prepared using iTaq SYBR Green Master Mix (Bio-Rad) and dispatched in reaction volumes of 25 μl containing 10 % (2.5 μl) of lysis supernatant. Primers to amplify the F-encoded *repE* gene (repE F & repE R) or the chromosome-encoded *dnaN* gene (dnaN F & dnaN R) were used at 300 nM (**Supplementary Table 2**). Amplification was quantified in real-time by detecting DNA-bound SYBR Green on the FAM channel of a CFX1000 instrument (Bio-Rad) and cycle thresholds (Cq) were detected using the Maestro software (Bio-Rad).

