## Supplementary material for "Single-cell evidence for plasmid addiction mediated by toxin-antitoxin systems": SupplementaryTables.docx

**Supplementary Table 1: Plasmids used in this study**

| **Plasmid** | **Properties** | **Reference** |
| --- | --- | --- |
| pNF03 | Mini-F *sopB-mNG* Δ*ccdAB cat* | This study |
| pNF03*ccd* | Mini-F *sopB-mNG* *ccdAB+ cat* | This study |
| pNF04 | Mini-F *sopB-mNG aphA I-SceI_CS_* | This study |
| pNF04*ccd* | pNF04 *ccdAB_F_* | This study |
| pNF04*vap* | pNF04 *vapBC_pINV_* | This study |
| pNF04*par* | pNF04 *parDE_RK2_* | This study |
| pNF04*doc* | pNF04 *phd-doc_P1_* | This study |
| pNF04*hok* | pNF04 *hok-sok_R1_* | This study |
| pNF04*hig* | pNF04 *higBA_Rts1_* | This study |
| pNF04*tac* | pNF04 *tacAT_p53668_226_* | This study |
| pNF06 | Mini-F Δ*sopABC* *aphA* | Jurenas, Fraikin et al., 2021, *mBio* |
| pNF06*ccd* | pNF06 *ccdAB_F_* | Jurenas, Fraikin et al., 2021, *mBio* |
| pSce | *ori_pSC101_* *cat* P*araBAD-SCE1* | This study |
| pJYB240 | Mini-F *sopB-mTurquoise2 cat* | J.Y. Bouet ; Guilhas et al., 2020, *Mol. Cell.* |
| pDL2655 | *ori_pSC101(ts)_* *cat* P*araBAD-SCE1* | C. Lesterlin |
| pUA66 | *ori_pSC101_* *aphA gfpmut2* | Zaslaver et al., 2006, *Nat. Meth.* |
| pCP20 | *ori_pSC101(ts)_ bla cat cI857* P*_L_-flp* | Cherepanov & Wackernagel, 1995, *Gene* |

**Supplementary Table 2: Oligonucleotides used in this study**

| **Name** | **Sequence** |
| --- | --- |
| ApaLI-ccdB F | ccccgtgcacgtctgctgtcagataaagtctcc |
| ApaLI-ccdB R | caaagtgcactggccaggggg |
| delccd F | atgtcaggctccgttatacac |
| delccd R | agcacacctctttttgacatac |
| NotI-pNF05bb F | cctatagcggccgcaaggcagttattggtgccc |
| HindIII-pNF05bb R | ccccaagcttcacatacgttccgccattcc |
| HindIII-KmR F | ccccaagcttatcgatgacgtcggaattgccagc |
| NotI-KmR R | cctatagcggccgcttactgtccctagtgcttgg |
| NotI-pNF04bb | cctatagcggccgcatgcgaaacgatcctcatcc |
| HindIII-pNF04bb R | ccccaagcttcacatacgttccgccattcc |
| 04Sce F | gttatccctatgtcccatcaggctttgc |
| 04Sce R | agggtaatctacacgaaggtttttgcgc |
| CmR-Sce F | gagtccaagctacctgtgacggaagatcac |
| CmR-Sce R | gcttgcatggccgcttatttcaggaaag |
| ori-Sce F | gaaataagcggccatgcaagctctagaggcatc |
| ori-Sce R | gtcacaggtagcttggactcctgttgatag |
| AatII-ccd F | ccccgacgtcttactaaaagccagataacagtatgcg |
| HindIII-ccd R | ccccaagctttatattccccagaacatcaggttaatg |
| AatII-vap F | ccccgacgtctgcccagcacatagtaattatcc |
| HindIII-vap R | ccccaagcttgaacaggtcagctccag |
| AatII-par F | ccccgacgtcattttcccgaccttaatgcg |
| HindIII-par R | ccccaagctttagcggctgaaatcagcc |
| AatII-doc F | ccccgacgtcttgtccggtcatttaagctgc |
| HindIII-doc R | ccccaagcttctcacgccatcaagaagcatc |
| AatII-hok F | ccccgacgtcgtgatgcggcaacaatc |
| HindIII-hok R | ccccaagcttcatccccgttcctgaag |
| AatII-hig F | ccccgacgtcttttgcaggctacgatctaccc |
| HindIII-hig R | ccccaagcttaatgctcttgcgcgttctgc |
| AatII-tac F | ccccgacgtctttgagtgcggttacgtcc |
| HindIII-tac R | ccccaagcttggatcggtcaactgatcc |
| repE F | gctgtatctgttcgttgacc |
| repE R | tcaatgcctgccgtatatcc |
| dnaN F | atcttcaatctgcacgctgg |
| dnaN R | aagagatcctcgacgttacc |

**Supplementary Table 3: Strains used in this study**

| **Strain** | **Genotype** | **Source** |
| --- | --- | --- |
| MG1655 | *E. coli* K-12 F- λ- *ilvG rfb-50 rph-1* | Lab Collection |
| FN042 | MG1655 *fhuA*::P*sulA-mTagBFP2-FRT* | Jurenas, Fraikin et al., 2021, *mBio* |
| *lon::tet* | MG1655 *lon*::ΔTn10 | Lab Collection |
| *gyrA462* | MG1655 *gyrA462* *zei*::Tn10 | Lab Collection |
| FN053 | FN042 *hupA-mCherry-FRT* | This study, P1vir LY119 x FN049 🡪 pCP20 |
| *recA* | MG1655 *recA_T233C_*-Tn10 | This study, P1vir LY542 x MG1655 |
| *recB* | MG1655 Δ*recB::FRT* | This study, P1vir JW2788 x MG1655 🡪 pCP20 |
| *ruvC* | MG1655 Δ*ruvC::FRT* | This study, P1vir JW1852 x MG1655 🡪 pCP20 |
| *lexA3* | MG1655 *lexA3*-Tn10 | Lab collection |
| *λ^+^* | MG1655 *λ^+^* | This study, lysogenization of MG1655 |
| *λ_imm21_^+^* | MG1655 *λ_imm21_^+^* | This study, lysogenization of MG1655 |
| *λ_imm434_^+^* | MG1655 *λ_imm434_^+^* | This study, lysogenization of MG1655 |
| *φ80^+^* | MG1655 *φ80^+^* | This study, lysogenization of MG1655 |
| *H-19B^+^* | MG1655 *H-19B^+^* | This study, lysogenization of MG1655 |
| LY542 | MG1655 *recA_T233C_*-Tn10 | C. Lesterlin |
| LY119 | MG1655 StrR *hupA-mCh-FRT-aphA-FRT* | C. Lesterlin |
| JW2788 | BW25113 Δ*recB::FRT-aphA-FRT* | KEIO collection |
| JW1852 | BW25113 Δ*ruvC::FRT-aphA-FRT* | KEIO collection |
| RH8000 | *E. coli* K-12 RK2^+^ | Lab Collection |
| BW27783 R1^+^ | BW25113 Δ*araEFG* ΔP*_araE_ ::*P*_CP8_*-*araE* R1^+^ | F. de la Cruz |

**Supplementary Legends**

**Supplementary Figure 1: Quantification of plasmid copy number by real-time polymerase chain reaction.** Chromosomal (*dnaN*) and plasmid-encoded (*repE*) fragments were amplified from crude DNA extracts. Data represents the mean and standard deviation of deltas in cycle of quantification (Cq) between *repE* and *dnaN*. ND : plasmid amplification could not be detected. ns : not significant (p = 0.2272, two-tailed t-test with Welch’s correction).

**Supplementary Figure 2: Quantification of I-SceI-induced plasmid curing by real-time polymerase chain reaction.** Cells were grown to exponential phase in MOPS medium containing 0.4 % glucose, 25 µg/ml kanamycin and 20 µg/ml chloramphenicol, then diluted 100x in MOPS medium containing 0.4 % arabinose and 20 µg/ml chloramphenicol. Samples were taken at indicated times for DNA extraction. Chromosomal (*dnaN*) and plasmid-encoded (*repE*) fragments were amplified from crude DNA extracts. Data represents three independent replicates.

**Supplementary Figure 3: Quantification of I-SceI-induced plasmid curing by time-lapse fluorescence microscopy.** FN053 cells transformed with pNF03 were grown in microfluidic chips under arabinose perfusion as in Figure 3A-B. Foci-free cells were counted at each 15-min timepoint.

**Supplementary Figure 4**: **Phage-induced viability loss under plasmid curing conditions.** Cells transformed with pSce and either pNF04 (EV) or pNF04ccd (*ccdAB*+) were grown to exponential phase in MOPS medium containing 0.4 % glucose, 25 µg/ml kanamycin and 20 µg/ml chloramphenicol, serially diluted and spotted on M9 plates containing 20 µg/ml chloramphenicol and either 0.4% glucose and 25 µg/ml kanamycin to promote plasmid retention, or 0.1% glucose and 0.3% arabinose to promote plasmid curing. Data represents the geometric mean and standard deviation of six independent experiments. a & b : ANOVA with post-hoc Tukey’s multiple comparison test. Note that data plotted for wt EV and wt *ccdAB*+ uses the same dataset as Figure 2B.

**Supplementary Figure 5: PSK-induced blebbing and lysis.** FN053 cells (MG1655 HU-mCherry) were transformed with pSce and pNF04 derivatives encoding the indicated TA systems, grown on MOPS medium containing 0.4 % glucose, 25 µg/ml kanamycin and 20 µg/ml chloramphenicol, loaded on microfluidic chips perfused with MOPS medium containing 0.4 % arabinose and 20 µg/ml chloramphenicol at time 0 to induce plasmid curing and imaged by time-lapse fluorescence microscopy every 15 min. Images show representative micrographs of plasmid-free segregants following I-SceI-mediated curing. SopB-mNeongreen is shown in green, the HU-mCherry fusion that localizes the chromosome is shown in red and the *sulA-bfp* fusion is shown in fire LUT. Each horizontally conjoined frames are taken 15 min apart. Scale bar is 1 µm. White arrows show loss of cytosolic content while blue arrows show blebbing.

**Supplementary Movies: Time-lapse fluorescence microscopy analysis of I-SceI-mediated curing of plasmids encoding various TA systems.** FN053 cells (MG1655 HU-mCherry, Movies 1-2, 5-10) or MG1655 cells lysogenized with lambda (Movies 3-4) were transformed with pSce and pNF04 derivatives encoding the indicated TA systems, grown on MOPS medium containing 0.4 % glucose, 25 µg/ml kanamycin and 20 µg/ml chloramphenicol, loaded on agarose pads (Movies 3-4) or microfluidic chips (Movies 1-2, 5-10) perfused with MOPS medium containing 0.4 % arabinose and 20 µg/ml chloramphenicol at time 0 to induce plasmid curing, then imaged by time-lapse fluorescence microscopy every 15 min. Scale bar is 5µm. **1:** Empty vector (EV) in microfluidic chip. **2:** *ccdAB* in microfluidic chip. **3:** Lambda lysogen with empty vector (EV) on agarose pad. **4:** Lambda lysogen with *ccdAB* on agarose pad. **5:** *parDE* in microfluidic chip. **6:** *hok-sok* in microfluidic chip. **7:** *higBA* in microfluidic chip. **8:** *vapBC* in microfluidic chip. **9:** *phd-doc* in microfluidic chip. **10:** *tacAT* in microfluidic chip
