## Supplementary figures and images for "Single-cell evidence for plasmid addiction mediated by toxin-antitoxin systems"

### FigS1.pdf

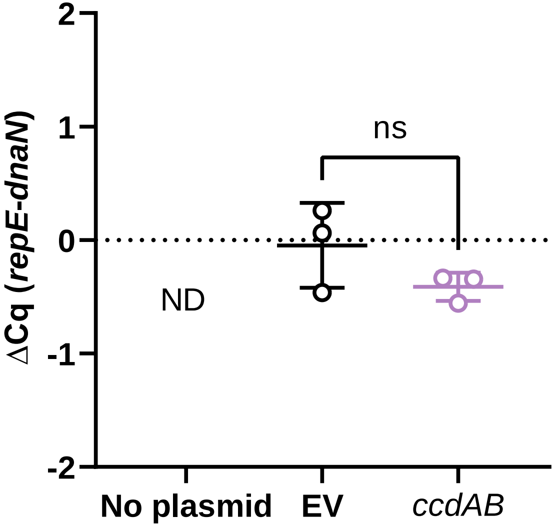

### FigS2.pdf

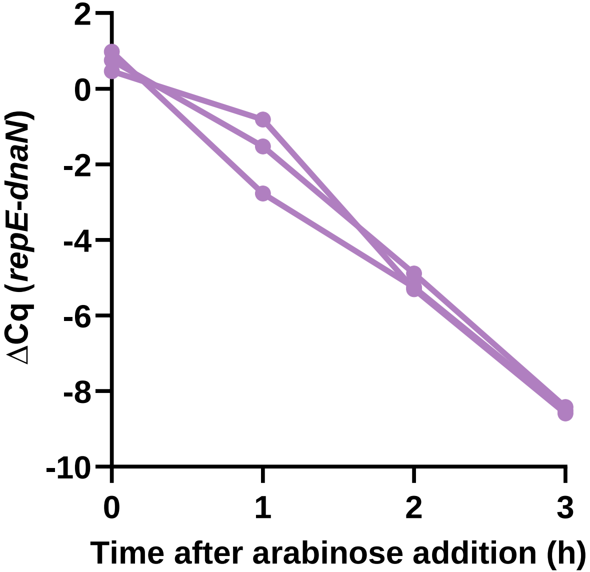

### FigS3.pdf

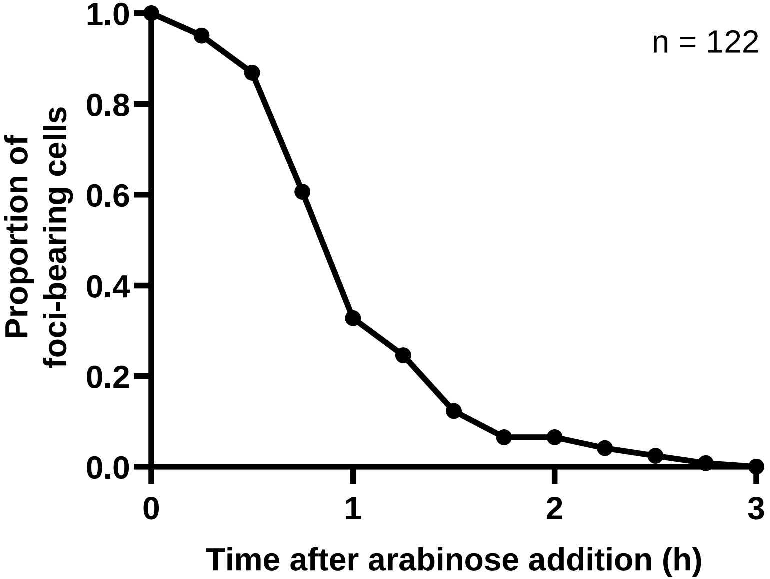

### FigS4.pdf

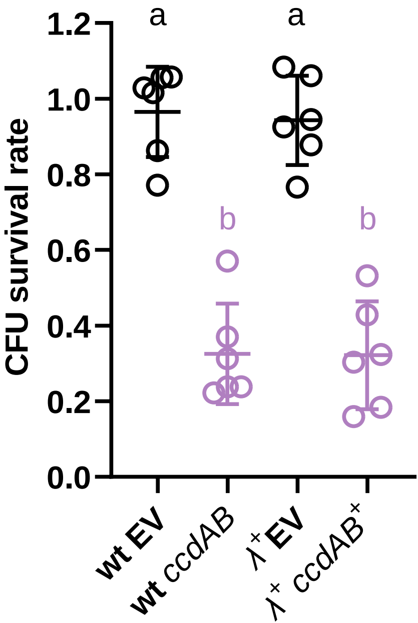

### FigS5.pdf

*phd-doc*$^{+}$

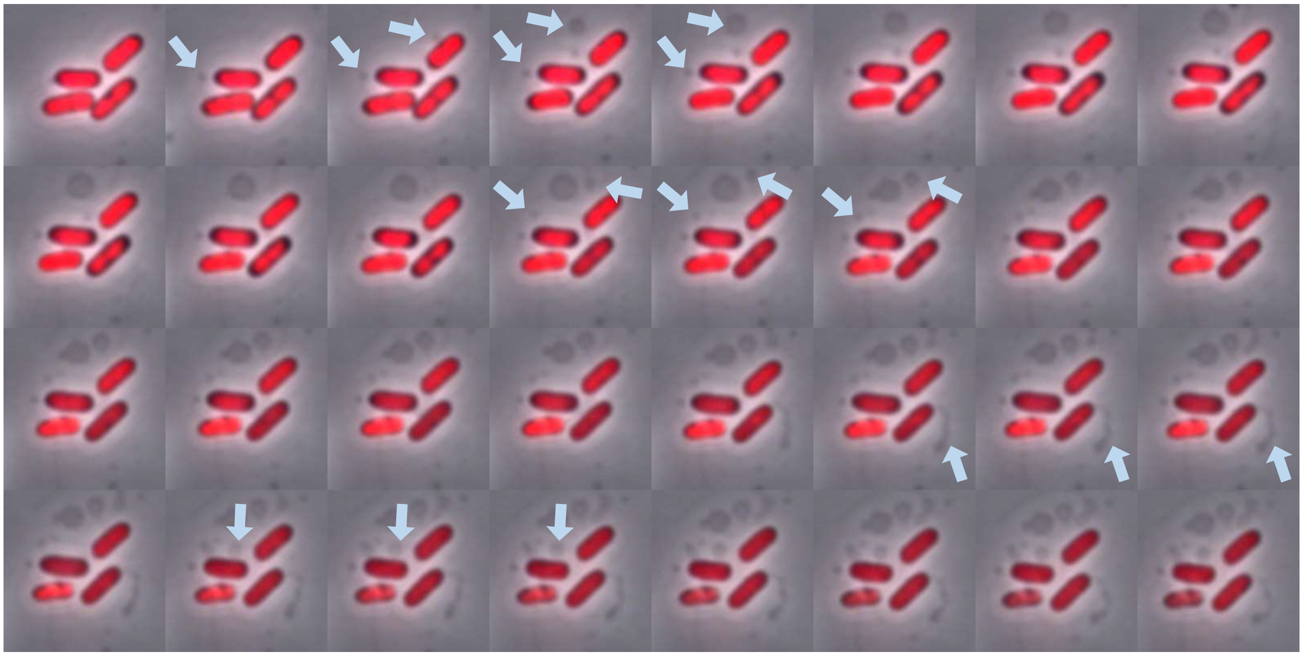

*hok-sok*$^{+}$

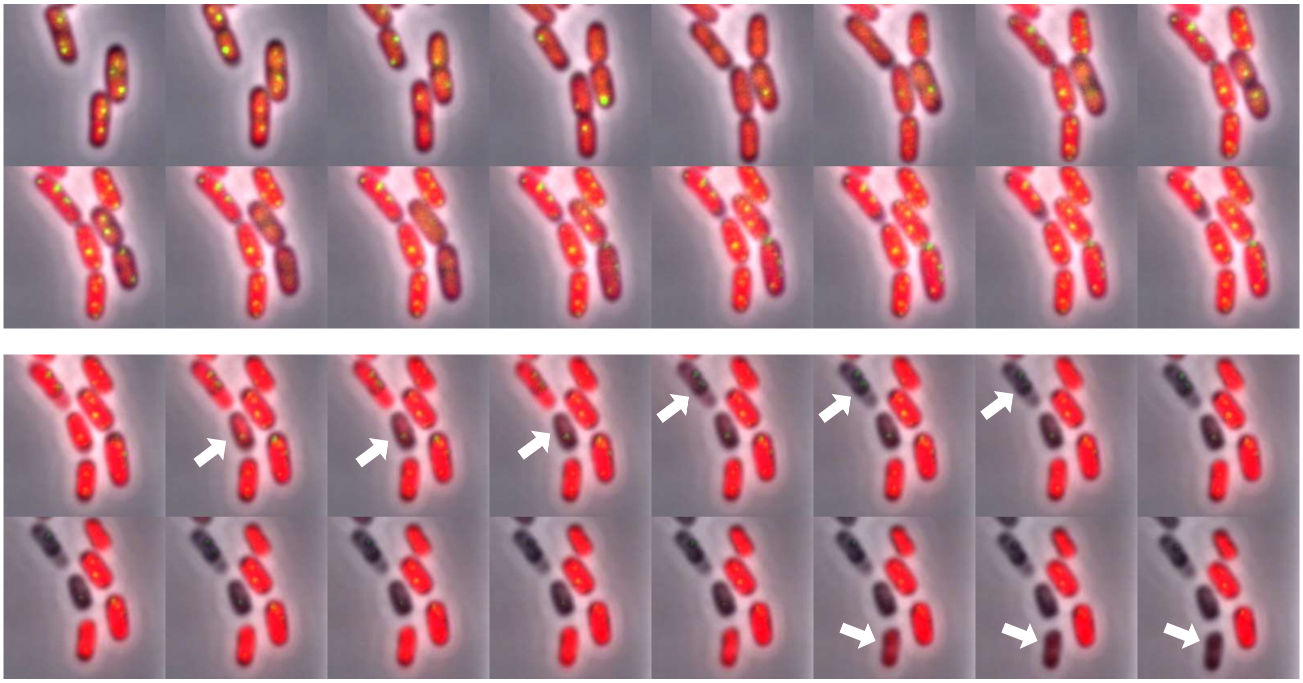

*tacAT*$^{+}$

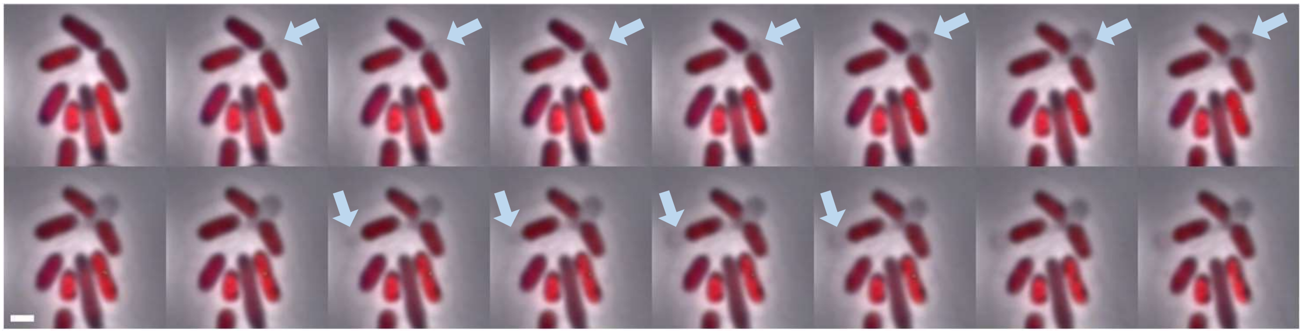
